# Circulating miR-126-3p is a mechanistic biomarker for knee osteoarthritis

**DOI:** 10.1101/2024.05.31.596603

**Authors:** Thomas G. Wilson, Madhu Baghel, Navdeep Kaur, Indrani Datta, Ian Loveless, Pratibha Potla, Devin Mendez, Logan Hansen, Kevin Baker, T. Sean Lynch, Vasilios Moutzouros, Jason Davis, Shabana Amanda Ali

## Abstract

As a chronic joint disease, osteoarthritis (OA) is a major contributor to pain and disability worldwide, and yet there are currently no validated soluble biomarkers or disease-modifying treatments. Since microRNAs are promising mechanistic biomarkers that can be therapeutically targeted, we aimed to prioritize reproducible circulating microRNAs in knee OA. We performed secondary analysis on two microRNA-sequencing datasets and found circulating miR-126-3p to be elevated in radiographic knee OA compared to non-OA individuals. This finding was validated in an independent cohort (N=145), where miR-126-3p showed an area under the receiver operating characteristic curve of 0.91 for distinguishing knee OA. Measuring miR-126-3p in six primary human knee OA tissues, subchondral bone, fat pad and synovium exhibited the highest levels, and cartilage the lowest. Following systemic miR-126-3p mimic treatment in a surgical mouse model of knee OA, we found reduced disease severity. Following miR-126-3p mimic treatment in human knee OA tissue explants, we found direct inhibition of genes associated with angiogenesis and indirect inhibition of genes associated with osteogenesis, adipogenesis, and synovitis. These findings suggest miR-126-3p becomes elevated during knee OA and mitigates disease processes to attenuate severity.

## Introduction

Osteoarthritis (OA) is a highly prevalent chronic joint disease that is estimated to reach up to 1 billion cases worldwide by 2050, with the knee being the most commonly affected joint^1^. Classically characterized by cartilage degradation^2^, current understanding presents OA as far more complex, with knee OA involving multiple other joint tissues including subchondral bone, synovium, fat, ligaments and meniscus^3^. There are presently no approved disease-modifying OA drugs (DMOADs), with treatments limited to symptom management before surgical interventions are ultimately indicated^4^. As such, there is a need to identify minimally-invasive molecular biomarkers that can be used to detect OA when opportunities for preventative interventions still exist^4^, and to better stratify individuals for recruitment to clinical trials evaluating DMOADs. MicroRNAs – small, non-coding RNA molecules – have emerged as a promising new class of biomarkers with the potential to meet this need^5^.

As biomarkers, microRNAs have numerous advantages^5^. First, they are accessible via minimally-invasive liquid biopsies, as compared to more invasive synovial fluid or tissue biopsies^6^. Second, their short length (20-24 nucleotides) and frequent encapsulation in microvesicles make them resistant to enzymatic degradation and more stable than other molecules^6^. Third, microRNAs can be reliably quantified such that small changes can be linked to disease outcomes^7^. Fourth, microRNAs show tissue-specific expression patterns, making it possible to map their role in complex diseases^8^. Fifth, microRNAs are known drivers of OA pathology, and their expression precedes phenotypic changes in tissues^9^. MicroRNAs are already in clinical use as biomarkers for musculoskeletal disorders such as osteoporosis^10^. Despite the potential of circulating microRNAs to serve as biomarkers of knee OA, a lack of reproducibility across microRNA profiling studies continues to be a hurdle to clinical translation.Beyond biomarkers, microRNAs are important epigenetic factors with mechanistic roles in musculoskeletal health and disease, and therefore represent promising therapeutic targets^11^. Produced in cells throughout the body, microRNAs are encoded in host genes and transcribed as primary stem loop structures (pri-microRNAs) that undergo enzymatic processing in the nucleus followed by transport to the cytoplasm to become mature microRNAs^12^. Primarily functioning to inhibit gene target expression through direct seed sequence binding, microRNAs are known to impact a variety of disease processes in OA, including inflammation, extracellular matrix dysregulation, and cell death and proliferation^9, 11, 13^. A notable therapeutic advantage of microRNAs, they can be readily modulated with small molecules in a targeted manner^14, 15^.

The gold standard for microRNA discovery is sequencing, which enables sensitive, specific, and high-throughput quantification of microRNAs in a given biospecimen^16^. To date there are only two published microRNA-sequencing studies that have evaluated circulating microRNAs between individuals with and without OA^17, 18^. The first study reported no significant differences in OA plasma extracellular vesicle microRNAs^17^, and the second study reported three differentially expressed (DE) microRNAs in OA serum, none of which were validated in subsequent experiments^18^. In the current study, we leveraged a customized microRNA-sequencing analysis pipeline^16^ to re-analyze these two datasets in search of reproducible circulating microRNAs associated with knee OA, and prioritized miR-126-3p. We then characterized miR-126-3p levels and potential mechanisms of action using both primary human knee OA tissues and a surgical mouse model of knee OA. Overall, our findings suggest miR-126-3p becomes elevated during knee OA as a mechanism to mitigate disease processes and attenuate OA severity.

## Results

### Circulating miR-126-3p is elevated in knee OA versus non-OA in independent cohorts

Taking an unbiased approach to identifying circulating microRNAs associated with knee OA versus non-OA, we leveraged two existing microRNA-sequencing datasets. These studies comprised Cohort 1 from Norway^17^ and Cohort 2 from France^18^ that previously reported zero and three DE microRNAs, respectively, none of which were subsequently validated. Leveraging the raw data from these studies, we performed secondary analysis using a customized microRNA-sequencing pipeline^16^ developed in previous studies^7, 19^ (**Figure 1A**). Specifically, we first defined knee OA based on Kellgren-Lawrence (KL) radiographic grade^20^, considering KL 3 or 4 as knee OA, and KL 0 as non-OA controls. These definitions were more stringent than those used in the original analyses, which for example, included KL 1 for controls^18^. Next, we performed a two-step read alignment, utilizing both miRBase v22.1 and the human reference genome (GRCh38), which captured additional microRNA reads that would otherwise not be aligned using a single reference database^16^. We then performed filtering to select microRNAs with a minimum of ten counts-per-million (CPM) in two or more samples and normalized the counts to total aligned sequences, consistent with previous studies^7, 19^. Differential expression analysis identified 23 microRNAs in OA versus non-OA individuals from Cohort 1 and 82 microRNAs from Cohort 2 at p < 0.1, with three DE microRNAs in common: miR-126-3p, miR-30c-2-3p, and miR-144-5p (**Figure 1A**). Amongst these three microRNAs, miR-126-3p exhibited the highest CPM (i.e., abundance), showed a consistent positive fold change in OA versus non-OA, and had the lowest p-value in both cohorts (**Figure 1B**). This data-driven discovery of circulating miR-126-3p in radiographic knee OA led us to prioritize it for further characterization.

**Figure 1.**
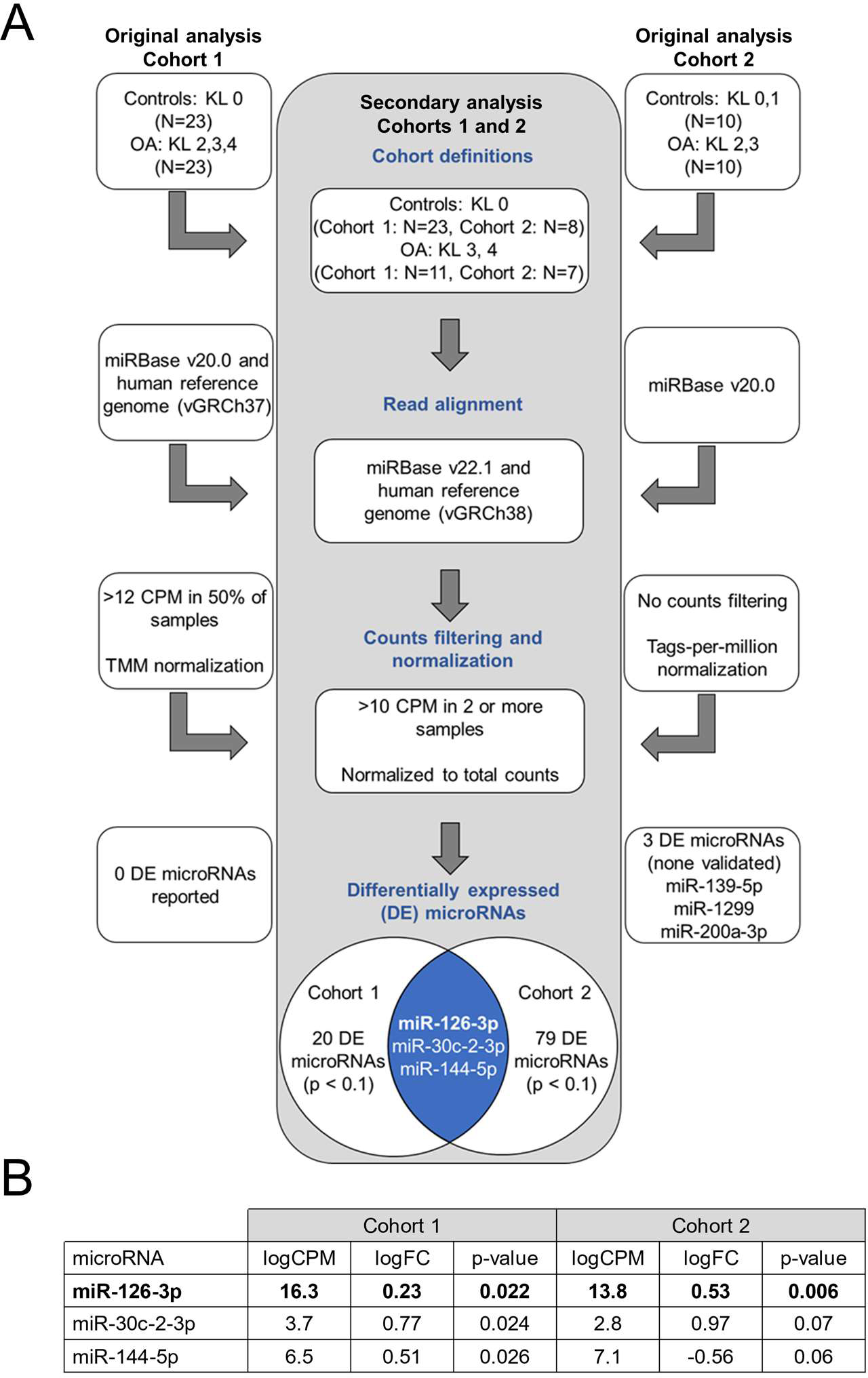
Circulating miR-126-3p is upregulated in knee OA versus non-OA in two independent microRNA-sequencing datasets. A) Overview of secondary analysis of two microRNA-sequencing datasets analyzed according to a customized analysis pipeline^16^. Left and right columns show details from the original analyses of Cohort 1^17^ and Cohort 2^18^, respectively, while the center column highlights our modifications and results. KL = Kellgren-Lawrence grade, CPM = counts-per-million, TMM = trimmed mean of m-values. B) Three differentially expressed microRNAs in knee OA versus non-OA were common in both datasets. LogCPM = log_2_ counts-per-million, logFC = log_2_ fold-change.

### Circulating miR-126-3p can accurately distinguish radiographic knee OA

To investigate miR-126-3p as a candidate biomarker for radiographic knee OA, we sought to define the patient population in which it is elevated using our Henry Ford Health (HFH) OA Cohort, a collection of primary human biofluids, tissues, and clinicodemographic data from consenting patients undergoing knee or hip arthroplasty or arthroscopy (**Supplemental Table 1**). After stratifying by joint (knee or hip) and KL grade, we measured miR-126-3p in plasma by real-time polymerase chain reaction (RT-PCR) and found that it was significantly elevated in KL ≥ 2 knee OA versus non-OA (controls with no evidence of OA), while notably, there was no increase in hip OA (**Figure 2A**). Since both microRNA levels^21^ and OA outcomes^22^ are influenced by age, sex, and body mass index (BMI), we next assessed the association of these variables with plasma miR-126-3p levels within knee OA patients using multiple linear regression analysis and found no association, whereas KL grade showed a significant positive association (**Figure 2B**). To assess the extent to which plasma miR-126-3p could be used to distinguish radiographic knee OA, we performed area under the receiver operating characteristic curve (AUC) analysis and found models including miR-126-3p had ‘excellent’ accuracy in distinguishing KL ≥ 2 knee OA from hip OA (AUC = 0.91, sensitivity = 0.91, specificity = 0.71; **Figure 2C**). This comparison was chosen since knee and hip OA groups exhibit similar clinicodemographic composition (**Supplemental Table 1**) but different plasma miR-126-3p levels (**Figure 2A**), where hip OA levels are similar to those in non-OA individuals. Based on its reproducibility across cohorts and accuracy in distinguishing KL ≥ 2 knee OA, our findings suggest circulating miR-126-3p is a promising candidate biomarker for radiographic knee OA.

**Figure 2.**
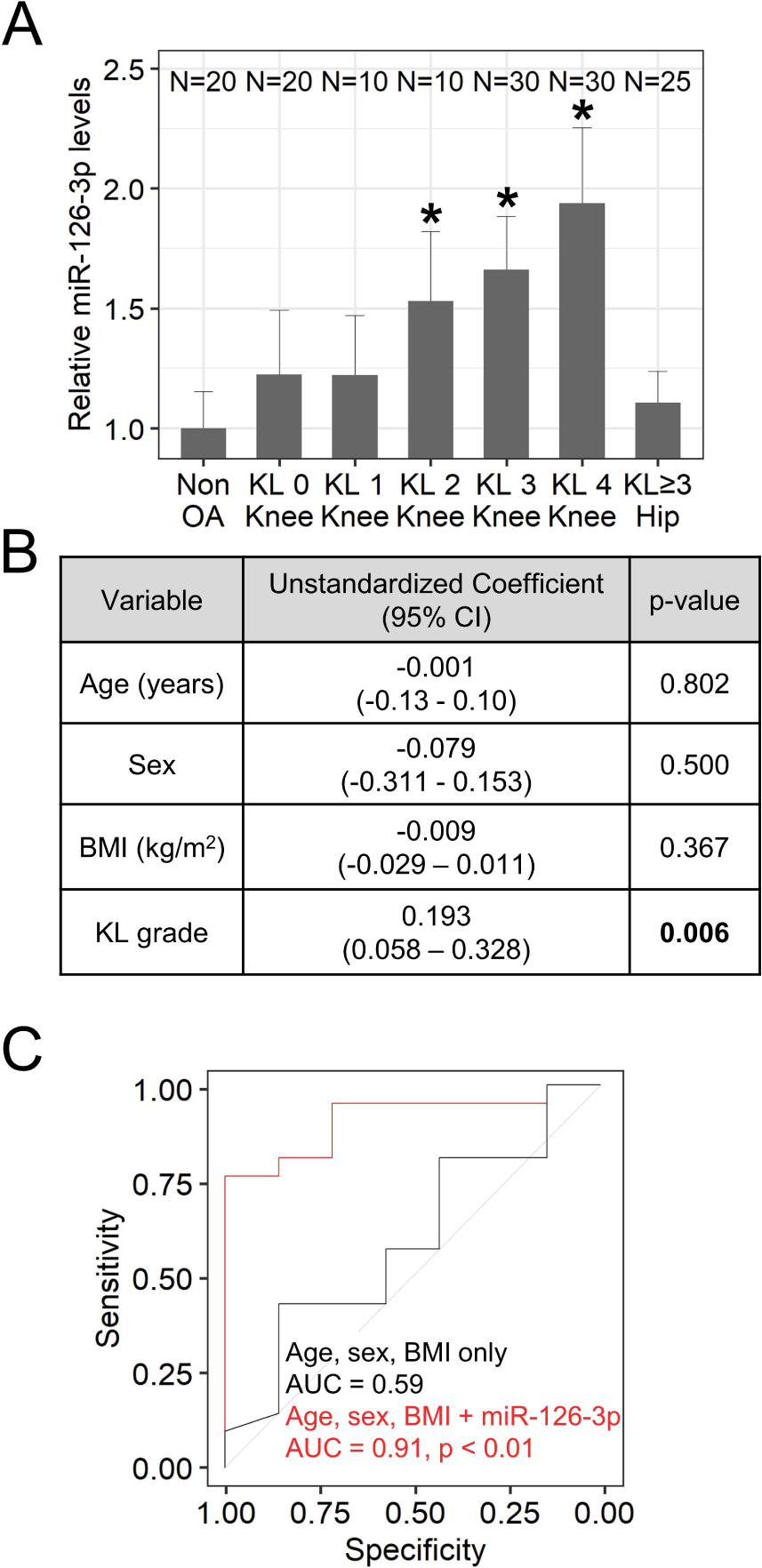
Circulating miR-126-3p distinguishes radiographic knee OA with excellent accuracy. A) Relative miR-126-3p expression in plasma samples collected from the Henry Ford Health (HFH) OA Cohort, stratified by joint and KL grade. Values are expressed as fold-change relative to average expression in non-OA controls. Error bars = 95% confidence interval, *p<0.05 versus non-OA. B) Multiple linear regression analysis assessing the association of each variable with plasma miR-126-3p levels in knee OA. BMI = body mass index, 95% CI = 95% confidence interval. C) Receiver operating characteristic curve analysis of a test cohort of HFH OA plasma samples displaying the accuracy of two models in distinguishing radiographic knee and hip OA. AUC = area under the receiver operating characteristic curve, p = versus age, sex, BMI only.

### Circulating miR-126-3p originates from knee OA fat pad and synovium

To assess the specificity of circulating miR-126-3p to knee OA, we measured mature miR-126-3p and its primary transcript (pri-mir-126) by RT-PCR in subchondral bone, infrapatellar fat pad (“fat pad”), synovium, anterior cruciate ligament, meniscus, and articular cartilage from individuals with OA undergoing total knee arthroplasty. First, we found mature miR-126-3p levels were highest in subchondral bone, fat pad and synovium as compared to cartilage, the tissue with the lowest average levels (**Figure 3A**). Second, we compared tissue miR-126-3p levels to plasma levels in matched samples and found significant positive correlations for subchondral bone, fat pad and synovium only (**Supplemental Table 2**). Third, we found pri-mir-126 expression was highest in fat pad and synovium, suggesting active transcription in these tissues, and relatively low in subchondral bone given its high levels of mature miR-126-3p (**Figure 3B**). Based on these findings, we hypothesized fat pad is a putative source of circulating miR-126-3p, while subchondral bone is a putative sink. We next measured secretion of mature miR-126-3p over time by each of the six knee OA tissues *ex vivo*. Conditioned media from tissue explants was collected after 24, 48, and 72 hours of culture, and miR-126-3p measured by RT-PCR (**Figure 3C**). We found miR-126-3p in fat pad conditioned media increased over time, whereas levels in subchondral bone conditioned media decreased, supporting our hypothesis of fat pad as a putative source (**Figure 3D**). We also found miR-126-3p in synovium conditioned media increased over time, suggesting it may be acting as a secondary source (given its lower levels) of miR-126-3p (**Figure 3D**). Taken together these data link circulating miR-126-3p to knee OA tissues, with fat pad and synovium as putative source tissues, and subchondral bone as a putative sink.

**Figure 3.**
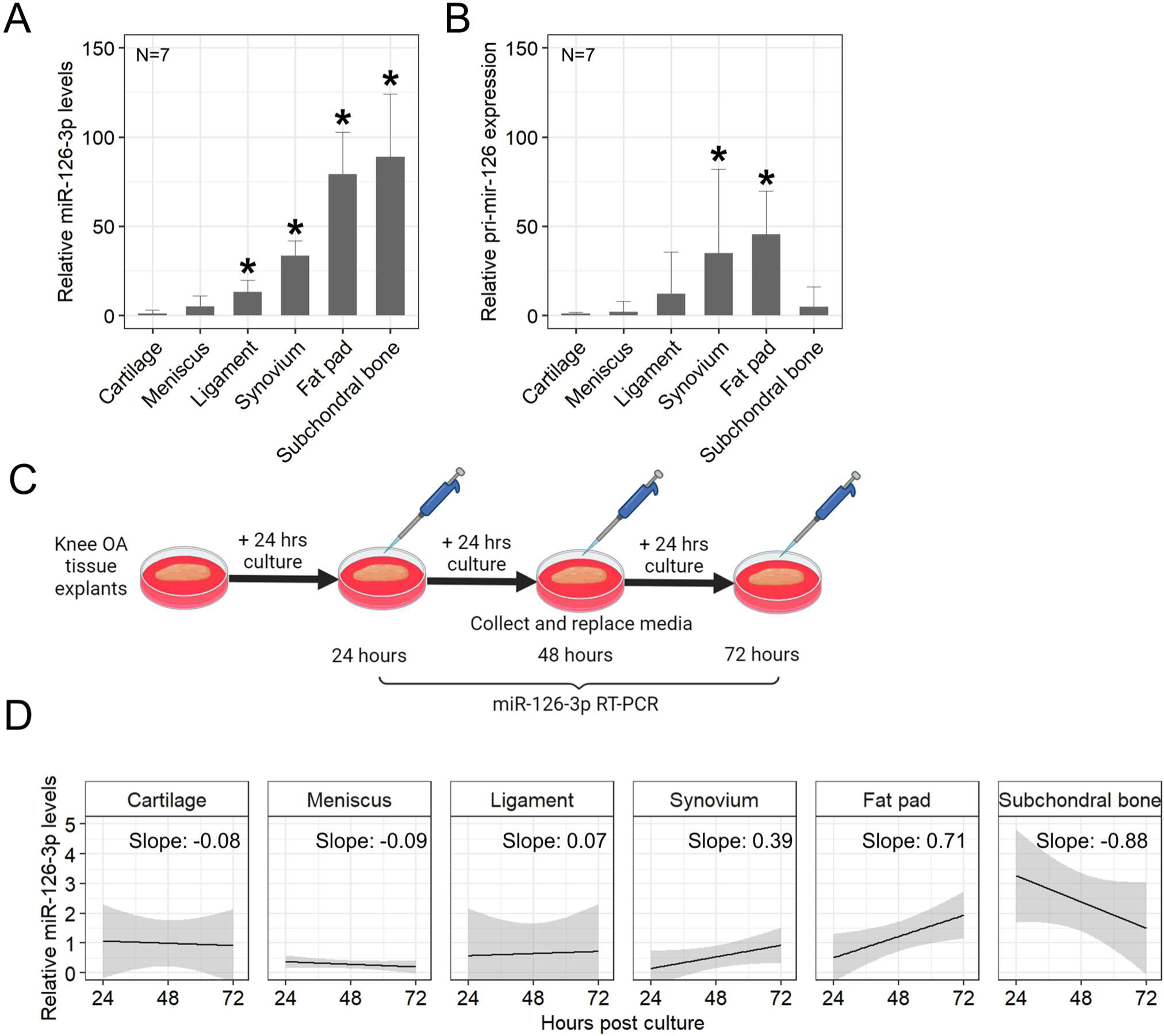
Knee OA fat pad and synovium are putative sources of miR-126-3p and subchondral bone a putative sink. A) Mature miR-126-3p levels and B) pri-mir-126 expression in six primary human knee OA tissues. Values represented as fold-change relative to average cartilage expression. Error bars = 95% confidence interval, *p<0.05 versus cartilage. C) Overview of experimental design for assessing miR-126-3p secretion over time. D) Secretion of miR-126-3p from six knee OA tissues. Values expressed as miR-126-3p fold-change relative to a reference microRNA (miR-24-3p) at each timepoint. Black line = fitted linear model, grey region = 95% confidence interval.

### miR-126-3p attenuates the severity of knee OA in a surgical mouse model

To investigate the effect of miR-126-3p on knee OA *in vivo*, we leveraged an established surgical mouse model comprising partial medial meniscectomy (PMX)^23, 24^ or sham surgery (**Figure 4A**). We compared miR-126-3p in plasma from PMX versus sham mice at four weeks post-surgery (16 weeks) and found an average 2.3-fold increase relative to pre-surgical levels (12 weeks; **Figure 4B**). This suggests knee OA is sufficient to induce elevated circulating miR-126-3p and supports the use of the PMX model for evaluating effects of miR-126-3p modulation on knee OA. Beginning four weeks post-operatively to approximate moderate knee OA^24^ (i.e., KL ≥ 2, when circulating miR-126-3p becomes elevated in humans; **Figure 2A**), weekly tail vein injections of 5 µg miR-126-3p mimic, inhibitor or negative control were delivered. After sacrifice (20 weeks), we performed OARSI histopathology scoring on Safranin-O stained sections of knee joints, where higher values reflect more cartilage damage^25^. We found miR-126-3p mimic treatment mitigated OA in PMX mice as compared to both miR-126-3p inhibitor and negative control treatments (**Figure 4C,D**). We also performed synovitis scoring (where higher values reflect more synovitis^26^) and similarly found better outcomes with miR-126-3p mimic in PMX mice as compared to miR-126-3p inhibitor (**Figure 4E,F**). There was no notable effect of these treatments in the sham groups for either outcome. Taken together, these data suggest that systemic delivery of miR-126-3p improves knee OA outcomes in a preclinical model.

**Figure 4.**
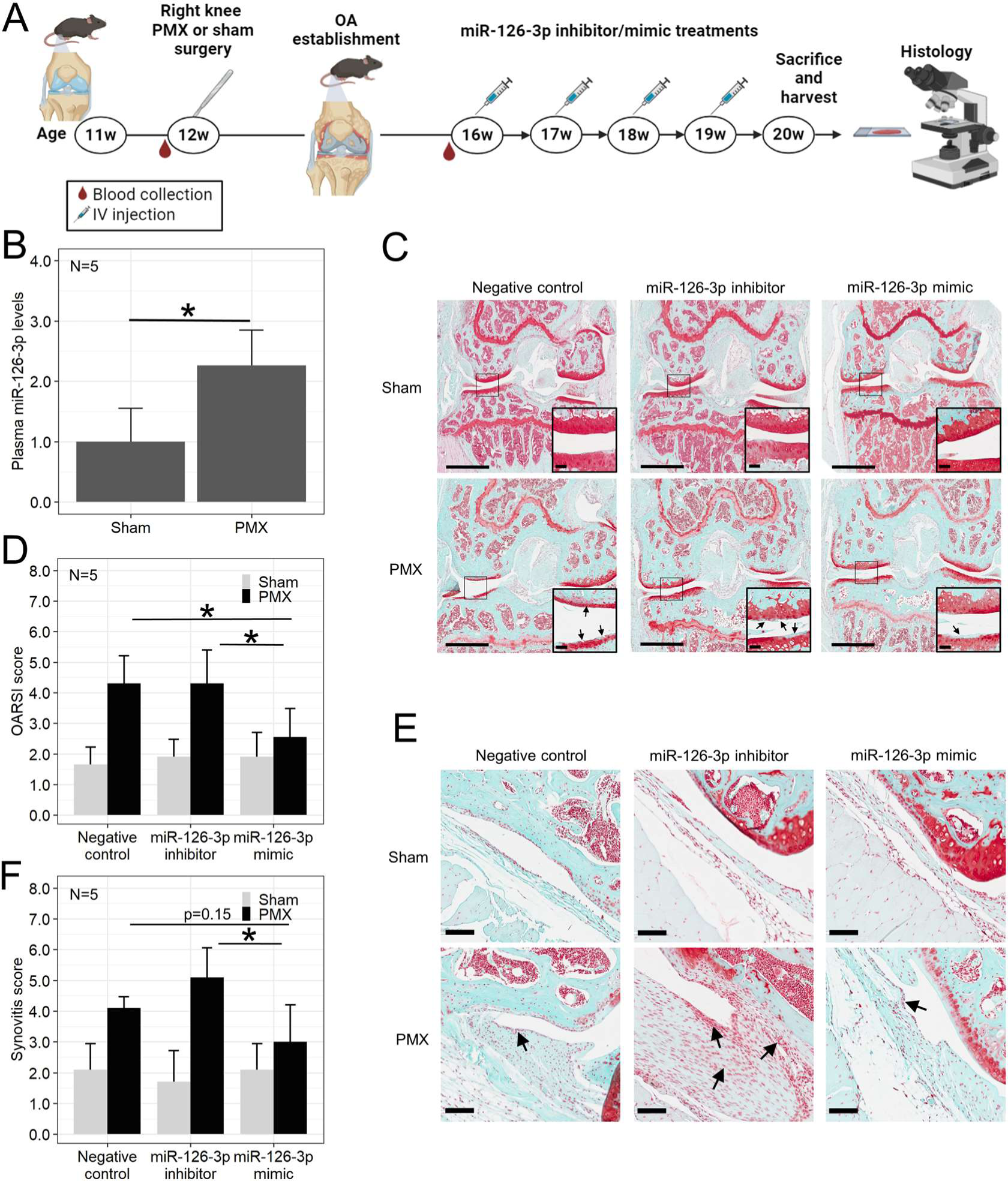
miR-126-3p improves outcomes in a surgical mouse model of knee OA. A) Overview of experimental design for mouse surgery, treatments, and endpoint. w = weeks old, PMX = partial medial meniscectomy, IV = intravenous. B) Plasma miR-126-3p levels in PMX versus sham mice at four weeks post-surgery (16w) relative to pre-surgery (12w). Error bars = 95% confidence interval, *p <0.05. C) Representative coronal sections of mouse right knee joint stained with Safranin-O. Scale bars = 1 mm. Insets show magnified regions (scale bars = 100 µm, arrows = examples of cartilage damage). D) OARSI scoring by blinded observers of PMX and sham mice following miR-126-3p treatments. Values are calculated as the average maximal quadrant scores for each treatment. Error bars = 95% confidence interval, *p<0.05. E) Representative sections of synovium from mouse right knee joint. Scale bars = 100 µm, arrows = examples of synovitis. F) Krenn synovitis scoring by blinded observers. Values are presented as average total synovitis scores for each treatment. Error bars = 95% confidence interval, *p<0.05.

### Direct and indirect gene regulation by miR-126-3p in knee OA

Based on previous literature in vascular systems, miR-126-3p is expressed by endothelial cells and functions to regulate angiogenesis^27^. We therefore identified validated direct gene targets of miR-126-3p associated with angiogenesis from the literature^27, 28, 29, 30, 31, 32, 33^ for assessment in the context of knee OA. Using RT-PCR, we first confirmed effective modulation of miR-126-3p in primary human knee OA subchondral bone, fat pad and synovium tissue explants following transfection with 100 nM miR-126-3p mimic (**Figure 5A,B**). We next measured changes in expression of Sprouty related EVH1 domain containing 1 (*SPRED1*), which is reported to inhibit angiogenesis^27, 28^, as well as A disintegrin and metalloproteinase domain 9 (*ADAM9*)^29^ and insulin receptor substrate 1 (*IRS1*)^30^, which are reported to enhance angiogenesis^31, 32, 33^. As expected of the inhibitory effect of microRNAs on their direct gene targets, we found reduced expression of all three genes with miR-126-3p mimic, except *IRS1* in synovium (p = 0.18; **Figure 5C-E**). While this leaves the net effect on angiogenesis unclear, we found that miR-126-3p lines blood vessels in each of the three tissue types, suggesting it may play a role in angiogenesis during knee OA (**Supplemental Figure 1**). Since angiogenesis is associated with knee OA outcomes^34^ we next investigated OA processes in each of subchondral bone, fat pad, and synovium that have been reported as secondary to angiogenesis^34, 35^. Following treatment with miR-126-3p mimic, we found decreased gene expression of osteocalcin (*OCN*), osterix (*OSX*) and runt-related transcription factor 2 (*RUNX2*) in subchondral bone (**Figure 5F**); decreased leptin (*LEP*), adiponectin (*ADIPOQ*) and a trend towards decreased CCAAT/enhancer-binding protein-alpha (*CEBPA*; p = 0.11) in fat pad (**Figure 5G**); and decreased interleukin 1 beta (*IL1b*), interleukin 6 (*IL6*) and tumor necrosis factor alpha (*TNFa*) in synovium (**Figure 5H**). Although likely through indirect mechanisms, these findings suggest miR-126-3p may mitigate knee OA by attenuating the osteogenesis associated with sclerosis and osteophytosis^36^, the adipogenesis associated with metabolic and signaling perturbations^37^, and the inflammatory response associated with synovitis^38^ (**Figure 6**).

**Figure 5.**
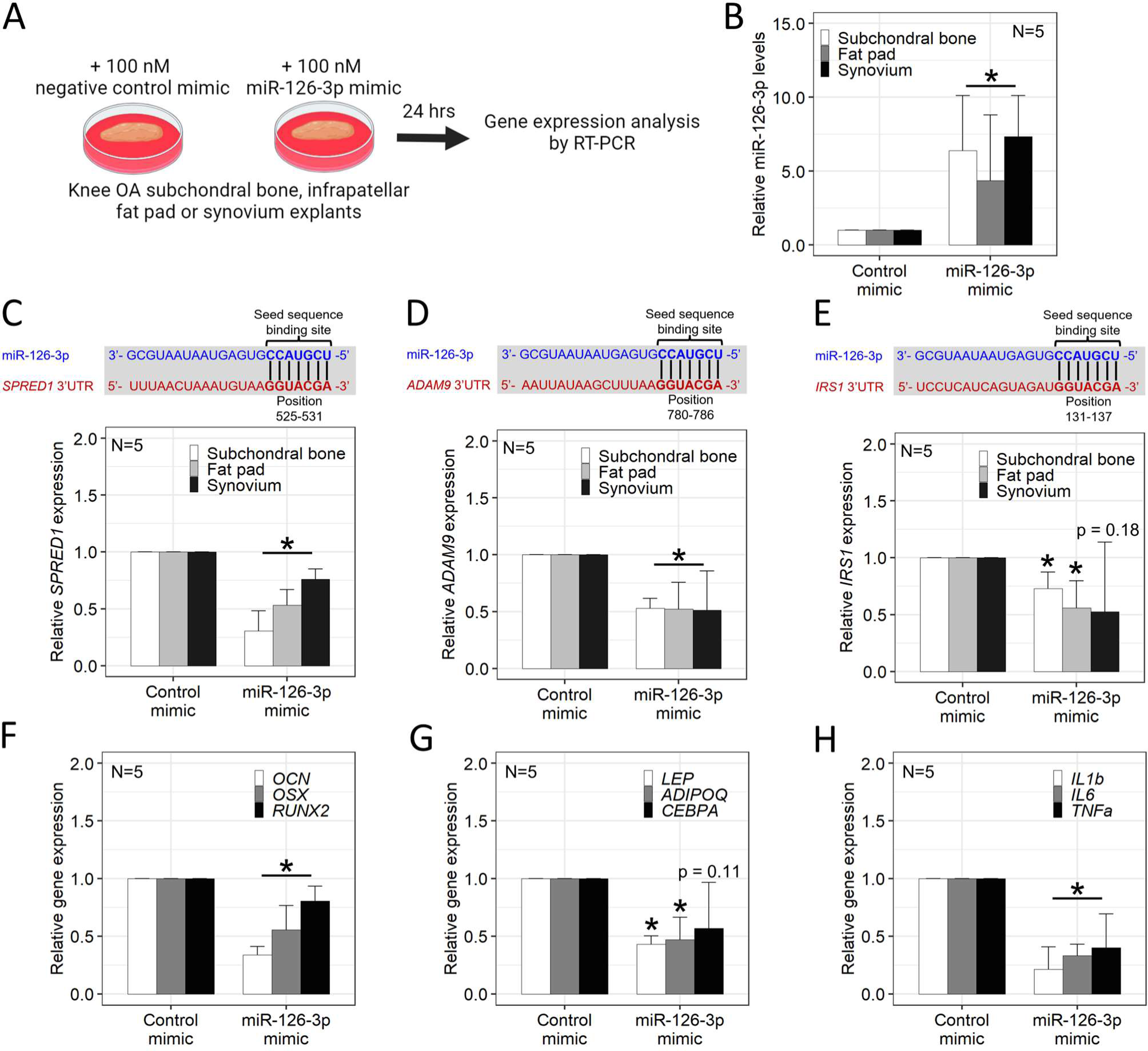
miR-126-3p modulates expression of direct and indirect gene targets in primary human knee OA tissues. A) Overview of experimental design for modulating miR-126-3p in knee OA tissues *ex vivo*. B) miR-126-3p levels in knee OA tissue explants following transfection with miR-126-3p mimic. C, D, E) Seed sequence binding site locations and gene expression changes in direct gene targets of miR-126-3p in knee OA tissue explants following transfection with miR-126-3p mimic. F, G, H) Gene expression changes in knee OA subchondral bone, fat pad and synovium tissue explants following transfection with miR-126-3p mimic. Values represent fold-changes relative to negative control mimic treated tissues. Error bars = 95% confidence interval, *p<0.05 versus negative control mimic. *SPRED1* = sprouty related EVH1 domain containing 1, *ADAM9* = A disintegrin and metalloproteinase domain 9, *IRS1 =* insulin receptor substrate 1, *OSX* = osterix, *OCN* = osteocalcin*, RUNX2* = runt-related transcription factor 2*, CEBPA* = CCAAT/enhancer-binding protein-alpha, *ADIPOQ* = adiponectin, *LEP* = leptin, *IL1b* = interleukin 1 beta, *IL6* = interleukin 6, *TNFa* = tumor necrosis factor alpha.

**Figure 6.**
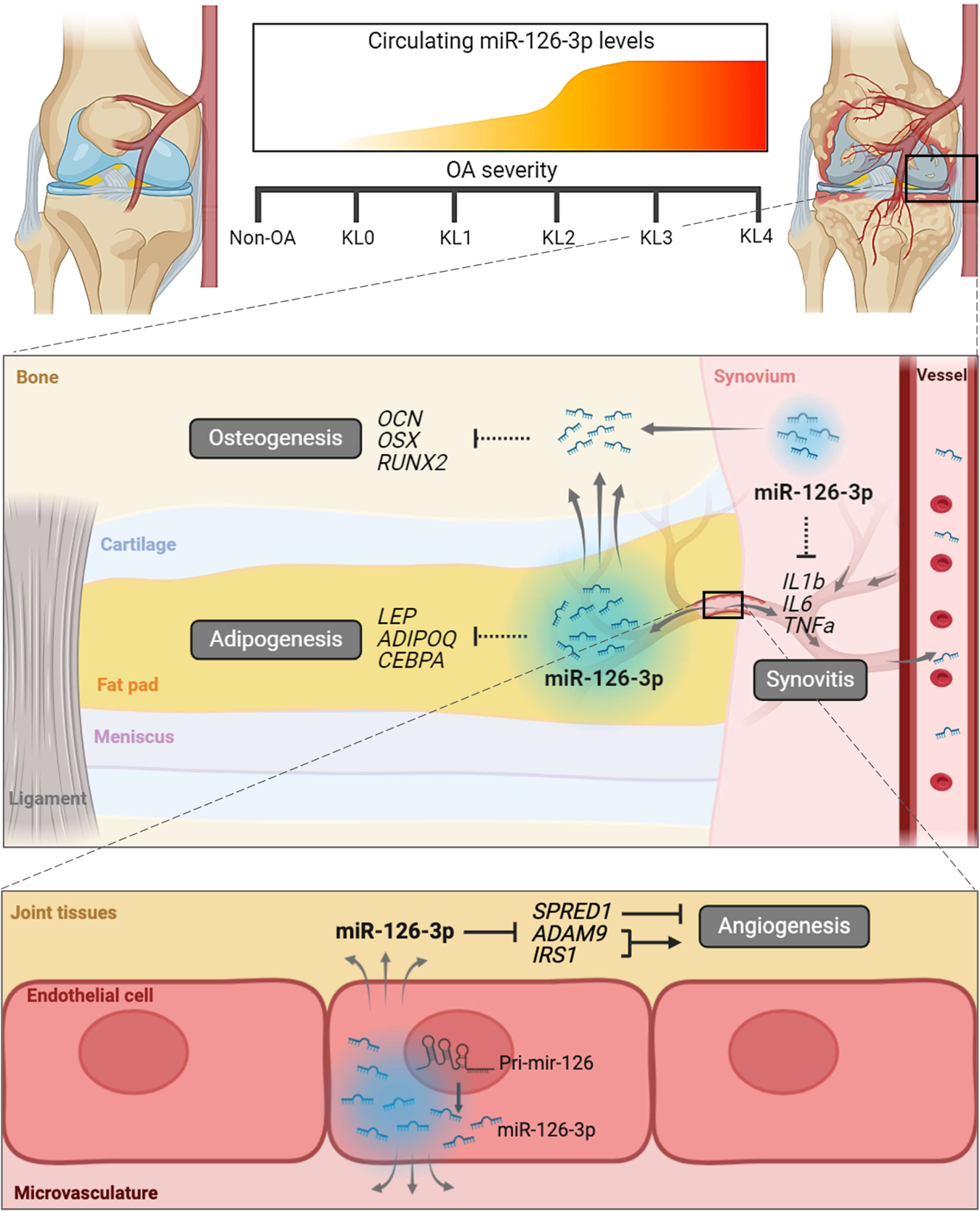
Proposed mechanism of action of miR-126-3p in knee OA. Circulating levels of miR-126-3p become elevated in KL ≥ 2 knee OA (top panel) via increased production of miR-126-3p (blue clouds) by endothelial cells in knee OA fat pad and synovium, which is secreted (grey arrows) into circulation though microvasculature and taken up by other knee OA tissues such as subchondral bone (middle panel). Within OA tissues, miR-126-3p indirectly reduces markers of osteogenesis in subchondral bone, adipogenesis in fat pad and synovitis in synovium (dashed black lines). These effects may be mediated by miR-126-3p regulation of angiogenesis through direct gene targets, including *SPRED1*, *ADAM9* and *IRS1* (solid black lines; bottom panel).

## Discussion

The objective of this study was to identify reproducible microRNAs as candidate mechanistic biomarkers for knee OA. With a data-driven approach, we leveraged a customized pipeline for microRNA-sequencing analysis^16^ and two previously published datasets^17, 18^ to prioritize circulating microRNAs in knee OA. We discovered and characterized miR-126-3p as a putative mechanistic biomarker for KL ≥ 2 knee OA, a stage considered to be moderate OA^39^, when opportunities for intervention to prevent or delay progression to end-stage disease still exist. This type of minimally-invasive and cost-effective biomarker could be useful for recruitment of more homogenous patient populations to clinical trials testing novel DMOADs or other OA therapies, including those focused on mitigating aberrant angiogenesis^40, 41^. In support of our finding, a previous study from Mexico reported increased levels of miR-126 in plasma from individuals with KL 2 and 3 knee OA^42^. Together with our HFH OA Cohort in the USA, this totals four independent datasets from four countries consistently showing elevation of miR-126-3p in knee OA versus non-OA controls, representing unprecedented reproducibility for a putative microRNA biomarker in the OA field to date^5^. Moreover, with respect to biofluids, two cohorts measured miR-126-3p in plasma^42^, one in serum^18^, and one in plasma extracellular vesicles^17^, and with respect to assays, two cohorts used sequencing^17, 18^, one used RT-PCR array^42^, and one used RT-PCR. Although low sample sizes limited statistical significance in the sequencing data after false-discovery-rate correction^43^, the diversity in biofluids and assays supports the robustness of our finding and points to the clinical utility of miR-126-3p as a biomarker wherein it could be measured in any readily accessible blood fraction using a common RT-PCR assay. Inclusion of miR-126-3p into composite biomarker profiles has proven useful for diseases like cancer^44^, suggesting the accuracy of our miR-126-3p model (AUC = 0.91) could be strengthened by including other emerging sensitive and specific biomarkers for knee OA (e.g., CRTAC1^5^) to improve identification of individuals with knee OA.

MicroRNAs are known to play important regulatory roles in biological processes that impact disease^9, 11^. Since microRNAs typically function to inhibit expression of their direct gene targets in a tissue-dependent manner^8^, we again undertook a data-driven approach to prioritizing knee OA tissues for further characterization of miR-126-3p. This proved to be an advantage over OA studies that focus on cartilage *a priori*, as our profiling of six different knee tissues revealed roles for subchondral bone, fat pad, and synovium, and relatively low levels of miR-126-3p in cartilage. A previous study exploring miR-126-3p in cartilage reported reduced levels in OA versus control and in old versus young cartilage^45^, whereas we found increased miR-126-3p levels in OA versus control plasma, and no association with age. Since we did not identify correlations between plasma and cartilage miR-126-3p levels in matched samples, the mechanisms governing miR-126-3p in cartilage may be distinct, particularly since cartilage is avascular and miR-126-3p is primarily expressed by endothelial cells^27, 46^. Our findings support the view of OA as a disease of the whole joint and put forth a role for tissue crosstalk with fat pad and synovium as potential source tissues of miR-126-3p and subchondral bone as a potential sink tissue. While our investigation into source and sink tissues was not exhaustive across the body, and more definitive experiments are required (e.g., tracing labeled microRNAs from source to sink^47^), data from our preclinical model suggest moderate knee OA is sufficient to produce elevated circulating levels of miR-126-3p. In support of fat pad as a putative source tissue, adipose-derived microRNAs are known to be a source of circulating microRNAs that can regulate genes in other tissues^48^. Similarly, in support of synovium as a putative source tissue and subchondral bone as a putative sink tissue, synovium-derived miR-126-3p-rich exosomes have been shown to attenuate subchondral bone phenotypes in an OA rat model^49^. In sum, we identified subchondral bone, fat pad and synovium as the most relevant tissues for understanding mechanisms of miR-126-3p in knee OA.

With evidence to support elevated circulating levels of miR-126-3p originating at least in part from knee OA tissues, we next used a preclinical model to investigate effects on knee OA outcomes and found systemic miR-126-3p mimic reduced cartilage damage and synovitis. These findings are consistent with the only other study exploring miR-126-3p in knee OA *in vivo,* reporting reduced osteophyte formation, cartilage degeneration and synovial inflammation in a surgical rat model following intra-articular delivery of exosomes carrying miR-126-3p^49^. Taken together this suggests that miR-126-3p has a pro-resolving effect in knee OA and may become elevated at KL ≥ 2 to mitigate disease processes. Though additional experiments are required to investigate the therapeutic potential of miR-126-3p, these data suggest there may be value in administering it earlier in the disease course (KL < 2) or at higher levels later in the disease course to improve knee OA outcomes. MicroRNAs are known to be promising therapeutic targets, with several clinical trials testing microRNA mimic-based treatments for conditions such as keloid disorders^50^, mesothelioma^51^ and advanced solid tumors^52^. Efforts are ongoing to overcome obstacles limiting microRNA-based therapies, including optimizing effective dosing and delivery methods, and minimizing unwanted off-target effects^14, 53^.

In terms of mechanisms through which miR-126-3p may be acting in knee OA, our data point to angiogenesis. In cancer and cardiovascular disease, miR-126-3p is known to regulate angiogenesis via direct targeting of *SPRED1*, *IRS1* and *ADAM9*, among others^30, 33, 54, 55^. In OA, angiogenesis is increased in multiple tissues including synovium, fat pad and subchondral bone^34, 56^, and is generally thought to be detrimental by promoting inflammation, pain and structural damage^34, 57^. The reduced gene expression in markers of osteogenesis, adipogenesis and synovitis we observed were through indirect mechanisms (i.e., not via direct miR-126-3p seed sequence binding), and therefore may be secondary effects of reduced angiogenesis^34, 35, 57^. Previous studies have shown that miR-126-3p can have both pro- and anti-angiogenic effects in a context-dependent manner^27, 58^. Among the three direct miR-126-3p gene targets we explored, one was anti-angiogenic (*SPRED1*^27, 28^) and two pro-angiogenic (*ADAM9*^31, 33^ and *IRS1*^32^), though all three showed reduced expression with miR-126-3p mimic, leaving the net effect on angiogenesis unclear in the absence of a functional assay. Future studies aimed at elucidating direct and indirect effects of miR-126-3p-mediated angiogenesis in knee OA, including potential effects on pain, are warranted to enhance our understanding of this mechanism. Additional studies are also required to assess joint-specific effects of miR-126-3p since our data show circulating levels are not elevated in hip OA, suggesting miR-126-3p-mediated mechanisms may be relevant to knee OA and not hip OA.

To our knowledge, this is the first study to identify a circulating microRNA that is consistently elevated in radiographic KL ≥ 2 knee OA versus non-OA individuals across four independent cohorts. We link this circulating microRNA to local knee tissues and show miR-126-3p exhibits a pro-resolving role in knee OA which may be directly mediated by angiogenesis, with secondary effects on osteogenesis, adipogenesis and synovitis. In sum, our findings suggest that miR-126-3p is a promising mechanistic biomarker for radiographic knee OA with therapeutic potential that merits further investigation.

## Methods

### HFH OA Cohort

All human biospecimens were obtained from our HFH OA Cohort. OA participants included individuals undergoing total joint arthroplasty or arthroscopy for the treatment of symptomatic knee or hip OA as assessed by an orthopedic surgeon. Severity was determined using radiographic KL grade^20^. Non-OA control participants consisted of individuals with no radiographic evidence of knee or hip OA, including some individuals undergoing hip femoroplasty, acetabuloplasty or labral repair. All biospecimens were collected, processed and stored according to our previously published protocols^59^. Knee OA tissues included subchondral bone, infrapatellar fat pad, synovium, anterior cruciate ligament, meniscus, and articular cartilage. Briefly, knee OA tissues were collected consistently by the same surgeons and dissected under sterile conditions within four hours of surgery, with subchondral bone and articular cartilage isolated from the anterior femoral condyle. Plasma was isolated from whole blood samples following centrifugation at 4000 rpm for 10 minutes at 4°C. Clinicodemographic data from each participant was captured and de-identified, including age, sex, BMI, race and comorbidities (**Supplemental Table 1**). The study protocol was approved by HFH Institutional Review Board (IRB #13995) and written informed consent was obtained from all participants prior to enrollment.

### Primary tissue explants

Primary human knee OA tissue explants (∼100 mg/well in a 24-well plate) were incubated in 500 µl Dulbecco’s Modified Eagle’s Medium (DMEM) supplemented with 1% penicillin-streptomycin (pen-strep) at 37°C, with 5% CO_2_. Secretion of microRNAs from knee OA tissue explants over time was assessed as outlined in **Figure 3C**. The collected media samples were centrifuged at 10000 x g for 5 minutes at 4°C to pellet cellular debris with the supernatant collected and stored at −80°C for further analysis.

### MicroRNA mimic and inhibitor transfection

As human and mouse miR-126-3p is homologous, we used the same miR-126-3p oligonucleotides for microRNA modulation in our primary human OA tissue explants and knee OA mouse model. Primary human knee OA tissue explants (∼100 mg/well in a 24-well plate) were cultured in 500 µl DMEM + 1% pen-strep. Tissues were transfected with 100nM miRIDIAN microRNA Human hsa-miR-126-3p Mimic (Dharmacon) or 100nM miRIDIAN microRNA Mimic Negative Control (Dharmacon) in combination with 2.5 µl/well DharmaFECT-1 Transfection Reagent (Dharmacon) at 37°C, as per the manufacturer’s recommendations. After 24 hours of incubation, tissues were collected, washed with sterile 1X phosphate buffered saline (PBS), flash frozen in liquid nitrogen and stored at −80°C for subsequent analyses. In mice, additional treatments of miRIDIAN microRNA Human hsa-miR-126-3p Hairpin Inhibitor (Dharmacon) and miRIDIAN microRNA Hairpin Inhibitor Negative Control (Dharmacon) were used as described below.

### Surgical mouse model of knee OA

Eleven-week-old male C57BL/6J mice were purchased from Jackson Laboratories and acclimatized in-house for one week. OA was induced via PMX, with the anterior half of the medial meniscus removed^23^. A concurrent group of mice were subjected to sham surgeries, with the medial meniscus left intact. Four weeks following surgeries, mice were randomized into treatment groups (N=5/group) and given weekly doses of 5 µg^15^ (in 1X PBS) miR-126-3p inhibitor, miR-126-3p mimic, or negative controls via tail vein injections for a total of four treatments. This *in vivo* experimental model is outlined in **Figure 4A**. For animal experiments, the negative control mimic and inhibitor treatments were combined (2.5 µg each in 1X PBS) to reduce excess animal burden, in accordance with the “three Rs” ethical guiding principles of animal research^60^. At endpoint, animals were euthanized and both hind limbs harvested. To assess the effects of knee OA on systemic miR-126-3p levels, blood samples were collected pre-surgery (12 weeks) and four weeks post-surgery (16 weeks). Plasma was isolated from whole blood samples following centrifugation at 4000 rpm for 10 minutes at 4°C. All animal experiments were approved by the HFH Institutional Animal Care and Use Committee (IACUC #1377).

### RNA extraction and quality assessment

For plasma and tissue culture conditioned media, RNA was extracted using the miRNeasy Serum/Plasma Advanced Kit (QIAGEN, Inc.) according to the manufacturer’s protocol. For tissue explants, RNA was isolated using our published protocols by phenol-chloroform extraction^59^. RNA concentration and quality were assessed using a NanoDrop 2000 spectrophotometer (Thermofisher Scientific).

### Real-time polymerase chain reaction (RT-PCR)

For microRNA quantification, reverse transcription was performed using the TaqMan microRNA Reverse Transcription Kit and TaqMan microRNA Assays (Applied Biosystems), according to the manufacturer’s instructions. For gene expression analysis, the High-Capacity cDNA Reverse Transcription kit (Applied Biosystems) was used to synthesize cDNA. RT-PCR was performed for both microRNA and genes using the QuantStudio 7 Pro Real-Time PCR System (Applied Biosystems) and TaqMan Gene Expression Assays (Applied Biosystems) or SYBR Green custom oligonucleotides (Sigma-Aldrich) as specified in **Supplemental Table 3**. *GAPDH* was used as the housekeeping gene for gene expression analyses and miR-24-3p as reference for microRNA quantification based on previous literature^15^. Results were analyzed using the delta-delta-Ct method^61^.

### Histological analyses

Harvested mouse knee joints were fixed in 10% neutral buffered formalin (NBF) at room temperature for 4 days and then decalcified using a 10% ethylenediaminetetraacetic acid (EDTA) solution (pH 7.6) at 4°C with agitation for 21 days. Following embedding, 5-micron coronal sections were placed on charged slides, deparaffinized and stained with 0.1% Safranin-O solution to assess OA severity. Changes in cartilage were assessed according to the OARSI histopathology guidelines for mice^25^. Knee sections were divided into quadrants (medial femur, medial tibia, lateral femur, lateral tibia) and graded from 0 (no damage) to 6 (> 75% cartilage erosion) by three independent blinded observers. The maximum quadrant scores by each observer for each section were averaged. Synovitis was assessed by the Krenn scoring system^26^. Values of 0 (normal) to 3 (severe) were assigned for enlargement of the synovial lining, stroma cell density and infiltration of inflammatory cells, then summed. Total scores from each blinded observer were averaged. MiR-126-3p localization was assessed in human tissues by *in situ* hybridization^62^. Knee OA subchondral bone, infrapatellar fat pad and synovium explants were fixed for 4 days in 10% NBF, after which bone samples were decalcified using a 10% EDTA solution, as described above. Fixed tissues were embedded and sectioned at 5-micron thickness, and mounted on charged slides. MiR-126-3p was then visualized using the SR-hsa-miR-126-3p-S1 miRNAscope probe (Advanced Cell Diagnostics) with the RNAScope 2.5 HD kit (Advanced Cell Diagnostics), according to the manufacturer’s instructions.

### Statistics

All data analysis was performed using R statistical software. Unless otherwise noted, statistical significance was determined using two-tailed Student’s T-test at p < 0.05 threshold. Differential expression analysis of microRNA-sequencing datasets was performed on normalized counts using a quasi-likelihood negative binomial regression model with trended dispersion. We trained a random forest classification model and assessed discrimination performance by AUC analysis. Input data were randomly split into training (70%) and testing (30%) cohorts. Sensitivity and specificity values were determined at a cut-point of 50% probability. Statistical significance between AUCs was determined by the DeLong test^63^. Multiple linear regression was performed for covariate analysis. Relationships between plasma and tissue microRNA levels were assessed by Pearson correlation.

## Acknowledgments

This study was funded in part by a grant awarded to SAA from the Arthritis National Research Foundation (ANRF 953971). We thank the authors of the original sequencing papers (Aae et al^17^, and Rousseau et al^18^) for providing raw sequencing data. We also thank participants of the Henry Ford Health OA Cohort for donating biospecimens.

## Author contributions

TGW, MB, and SAA were involved in study conception and design. TGW, MB and NK performed experimental work and data acquisition. PP, ID and IL were involved in secondary analysis of the sequencing datasets. LH, DM, KB, TSL, VM and JD contributed to the HFH OA Cohort biospecimens and data. TGW, MB and SAA performed data analysis, interpretation, and drafted the manuscript. All authors contributed to manuscript revisions and approved the final version.

## Supplemental Material

**Supplemental Figure 1.**
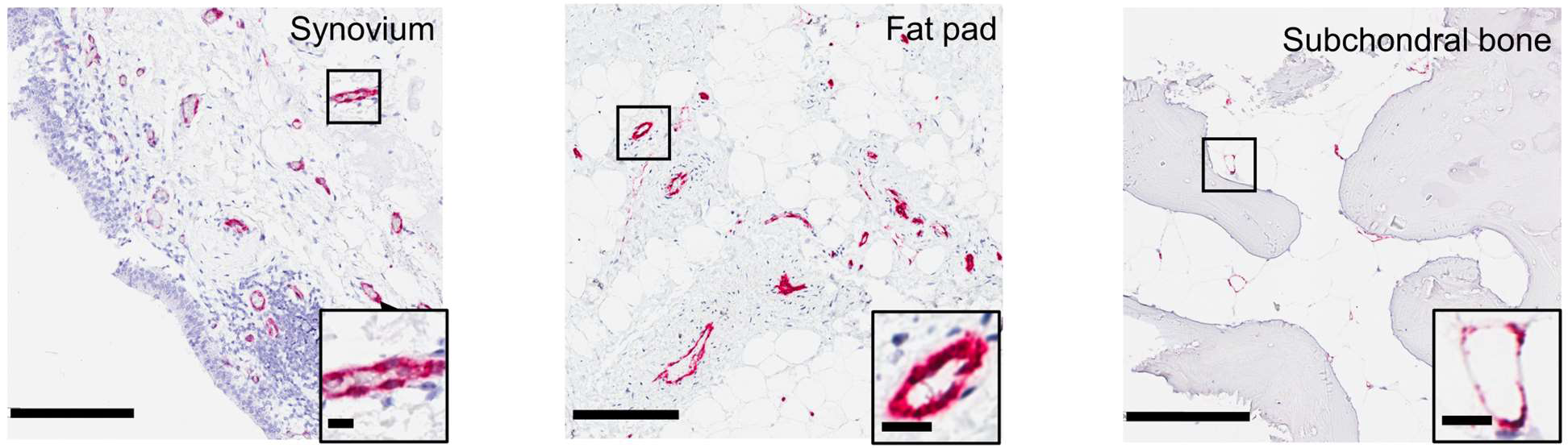
Representative images depicting localization of miR-126-3p in primary human knee OA tissues by *in situ* hybridization. Scale bars = 200 µm, red = miR-126-3p staining. Insets show magnified blood vessels (scale bars = 15 µm).

**Supplemental Table 1.** Key clinicodemographic variables from the Henry Ford Health OA Cohort. Top comorbidity = the most commonly reported comorbidity for each group. Significance between groups assessed by one-way ANOVA for continuous variables and Chi-square test for categorical variables. BMI = body mass index, SD = standard deviation, Hyp = hypertension.

|  | Non<br>OA | KL 0<br>Knee | KL 1<br>Knee | KL 2<br>Knee | KL 3<br>Knee | KL 4<br>Knee | KL ≥ 3<br>Hip | p-value |
| --- | --- | --- | --- | --- | --- | --- | --- | --- |
| N | 20 | 20 | 10 | 10 | 30 | 30 | 25 | - |
| Age (years)<br>Mean (SD) | 35.2<br>(12.3) | 27.9<br>(7.7) | 46.9<br>(10.1) | 56.2<br>(13.0) | 66.9<br>(7.3) | 67.4<br>(7.9) | 64.0<br>(9.6) | <0.01 |
| Sex (female)<br>N (%) | 17<br>(85.0) | 11<br>(55.0) | 3<br>(30.0) | 8<br>(80.0) | 20<br>(66.7) | 16<br>(53.3) | 13<br>(52.0) | 0.1 |
| BMI (kg/m<sup>2</sup>)<br>Mean (SD) | 25.8<br>(4.1) | 29.3<br>(8.1) | 29.5<br>(5.3) | 30.4<br>(4.2) | 32.4<br>(4.6) | 32.5<br>(5.4) | 29.7<br>(4.3) | <0.01 |
| Race (white)<br>N (%) | 20<br>(100) | 17<br>(85.0) | 8<br>(80.0) | 9<br>(90.0) | 25<br>(83.3) | 19<br>(63.3) | 23<br>(92.0) | 0.5 |
| Top comorbidity<br>(%) | Asthma<br>(10.0) | Asthma<br>(15.0) | Asthma<br>(30.0) | Hyp<br>(60.0) | Hyp<br>(66.0) | Hyp<br>(53.3) | Hyp<br>(52.0) | - |

**Supplemental Table 2.** Pearson correlation analysis between miR-126-3p levels in matched tissue and plasma samples from knee OA individuals. R = correlation coefficient.

| Tissue (N=18) | R | p-value |
| --- | --- | --- |
| Subchondral bone | 0.49 | **0.04** |
| Fat pad | 0.55 | **0.02** |
| Synovium | 0.59 | **0.01** |
| Ligament | -0.03 | 0.91 |
| Meniscus | 0.19 | 0.44 |
| Cartilage | -0.10 | 0.70 |

**Supplemental Table 3.**
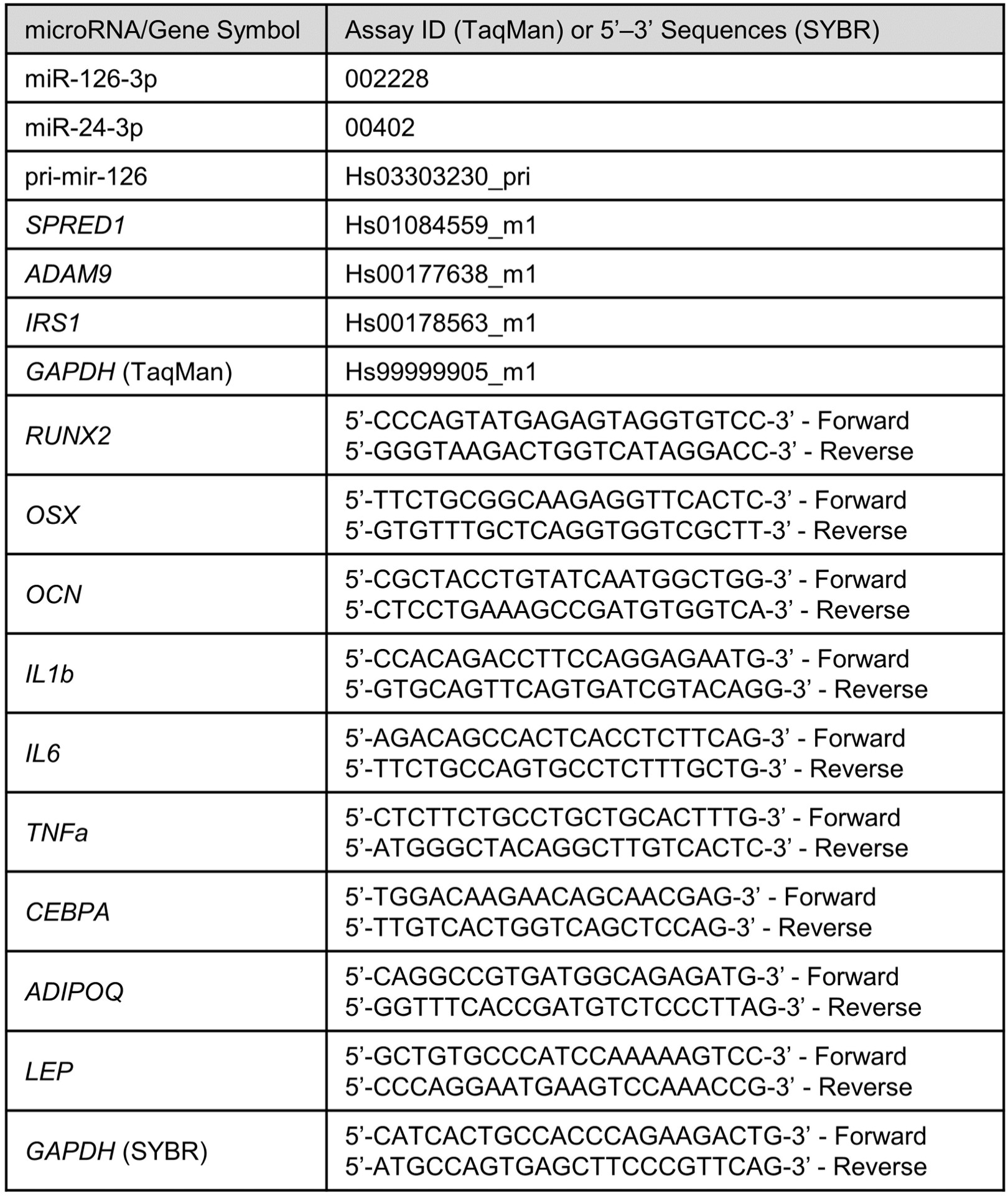
List of primers used for RT-PCR experiments.

